# Source range phylogenetic community structure can predict the outcome of avian introductions

**DOI:** 10.1101/2021.05.17.444023

**Authors:** Brian S. Maitner, Daniel S. Park, Brian J Enquist, Katrina M Dlugosch

## Abstract

Competing phylogenetic models have been proposed to explain the success of species introduced to other communities. Here, we present a study predicting the establishment success of birds introduced to Florida, Hawaii, and New Zealand using several alternative models, considering species’ phylogenetic relatedness to source and recipient range taxa, propagule pressure, and traits. We find consistent support for the predictive ability of source region phylogenetic structure. However, we find that the effects of recipient region phylogenetic structure vary in sign and magnitude depending on inclusion of source region phylogenetic structure, delineation of the recipient species pool, and the use of phylogenetic correction in the models. We argue that tests of alternative phylogenetic hypotheses including the both source and recipient community phylogenetic structure, as well as important covariates such as propagule pressure, are likely to be critical for identifying general phylogenetic patterns in introduction success, predicting future invasions, and for stimulating further exploration of the underlying mechanisms of invasibility.

## Introduction

Understanding why some species that have been introduced to a new location are able to establish self-sustaining populations while others fail to do so is of critical importance for preventing the spread of invasive species; increasing the success of deliberate introductions (e.g. reintroductions, assisted migrations, rewilding); and for understanding the community assembly process. With the growing availability of phylogenetic data (e.g. Jetz et al. 2012, Faurby and Svenning 2015, Smith and Brown 2018), there has been an increase of interest in methods that use phylogenetic data to try to predict which introduced species are more likely to become established (Strauss et al. 2006, Maitner et al. 2012, Park and Potter 2013, Ma et al. 2016).

Most uses of phylogenetic data to predict the success of introduced species rely on the assumption that closely-related species are more similar in both their competitive niches and environmental requirements (Darwin 1859, Webb 2000, Wiens et al. 2010). From this assumption, two competing hypotheses have been posited: the Competition-Relatedness Hypothesis (also known as Darwin’s Naturalization Hypothesis; Rejmánek 1996) and the Environmental Filtering Hypothesis (also known as the pre-adaptation hypothesis; Ricciardi and Mottiar 2006). The competition-relatedness hypothesis assumes that because niche differences likely increase with phylogenetic distance, competition will generally be more intense among closely-related species, leading to competitive exclusion (Darwin 1859). Conversely, the environmental filtering hypothesis assumes that introduced species closely-related to natives may generally be more likely to establish because they are adapted to similar environments (Darwin 1859). In summary, the competition-relatedness hypothesis predicts that introduced species that are closely-related to the recipient community will be *less likely* to successfully establish (due to competitive exclusion from close relatives; Table 1) while the environmental filtering hypothesis predicts that these species will be *more likely* to successfully establish (due to shared environmental tolerances with the recipient community; Table 1; Darwin 1859).

**Table 1.** Summary of phylogenetic hypotheses of establishment success. Hypotheses highlighted in blue predict that being closely-related to a community will increase establishment success, while those highlighted in red predict that being *distantly*-related to a community will increase establishment success.

|  | Hypothesis | Phylogenetic Predictions | Hypothesized Mechanisms | Citations |
| --- | --- | --- | --- | --- |
| Recipient Range | Environmental Filtering (EFH, Pre-Adaptation) | Species closely-related to recipient community are more successful | Shared environmental tolerances<br>Shared mutualists | Darwin 1859<br>Ricciardi and Mottiar 2006 |
|  | Competition-Relatedness (CRH, Darwin's Naturalization) | Species distantly-related to recipient community are more successful | Reduced competitive exclusion<br>Avoid shared natural enemies | Darwin 1859<br>Rejmánek 1996 |
| Source Range | Evolutionary Imbalance (EIH, Biogeographic Superiority) | Species from regions with many/close relatives are more successful | Increased phenotypic optimization due to high competition/apparent competition<br>Longer exposure to selection in a given environment<br>Competitive superiority | Darwin 1859<br>Fridley and Sax 2013 |
|  | Universal Trade-Off (UTH) | Success is independent of source community phylogenetic structure | Universal constraints limit species to a common trade-off surface | Niklas 1997<br>Marks and Lechowicz 2006<br>Tilman 2011 |
|  | Competitive Constraint (CCH) | Species from regions with few/distant relatives are more successful | Competition and/or apparent competition constrain adaptation | de Mazancourt et al. 2008<br>Meyer and Kassen 2007<br>Wilson 2014 |

Both the competition-relatedness hypothesis and the environmental filtering hypothesis focus solely on phylogenetic similarity between an introduced species and the recipient community. This makes the implicit assumption that the evolutionary context of the source region does not make a substantial contribution to establishment success. However, Maitner *et al. (2021)* highlight three additional hypotheses which make predictions about the establishment success of species based on their phylogenetic relationships with taxa in their native source regions: the Evolutionary Imbalance Hypothesis (Darwin 1859, Fridley and Sax 2013), the Competitive Constraint Hypothesis (Meyer and Kassen 2007, de Mazancourt et al. 2008, Wilson 2014), and the Universal Trade-Off Hypothesis (Niklas 1997, Marks and Lechowicz 2006, Tilman 2011). The Evolutionary Imbalance Hypothesis states that phenotypic optimization will be maximized in regions characterized by intense competition (i.e. many close relatives, if competition intensity declines with phylogenetic distance) that have experienced similar environmental conditions for longer periods of time (Fridley and Sax 2013). The Competitive Constraint Hypothesis proposes that species originating in regions containing many competitors (or apparent competitors) will be at a competitive disadvantage, as they may have experienced reduced population sizes leading to fewer beneficial mutations and reduced efficacy of selection (Meyer and Kassen 2007, Wilson 2014) and may have been constrained in their evolutionary trajectories by character displacement (de Mazancourt et al. 2008). The universal trade-off hypothesis states that potential competitors are subject to the same set of evolutionary trade-offs regardless of their source region, and as such will have similar species average fitnesses (Tilman 2011). The evolutionary imbalance hypothesis thus predicts that species originating from regions with many close relatives will be relatively successful invaders due to a greater phenotypic optimization (Table 1). Conversely, the competitive constraint hypothesis predicts that species originating from regions containing many close relatives will be relatively *un*successful invaders, as their evolutionary trajectories may have been constrained by strong competition. Finally, the universal trade-off hypothesis predicts that introduction success should be unrelated to source region phylogenetic structure, as species from different regions should be roughly equivalent in relative fitness.

Previous studies examining the predictive power of phylogenetic structure on establishment success have largely focused on the mean (or minimum) distance between an introduced species and the recipient community (e.g. Strauss et al. 2006, Maitner et al. 2012, Park and Potter 2013, 2015a, b, Marx et al. 2016, Ma et al. 2016). However, focusing solely on the mean (or minimum) distance between species ignores much of the information present within the distributions of phylogenetic distances which may be captured by the higher moments of the distributions (Fig. 1; Maitner et al. 2021). In situations where we expect being closely-related to a community to be beneficial (i.e. the environmental filtering and evolutionary imbalance hypotheses), we also predict that: (1) species with lower variance in phylogenetic distance should be more successful, as these species are distantly-related to the entire community; (2) species with a negative kurtosis will be more successful, as this reflects relatively few, closely-related species; and (3) species with a higher kurtosis with be more successful, as these species will be basal taxa that are similarly distantly-related to the rest of the community. Where we expect being distantly-related to a community to be beneficial (i.e. the competition-relatedness and competitive constraint hypotheses), the predictions are the opposite. Finally, in the case of the universal trade-off hypothesis, we expect these to be unrelated to establishment success.

**Figure 1.**
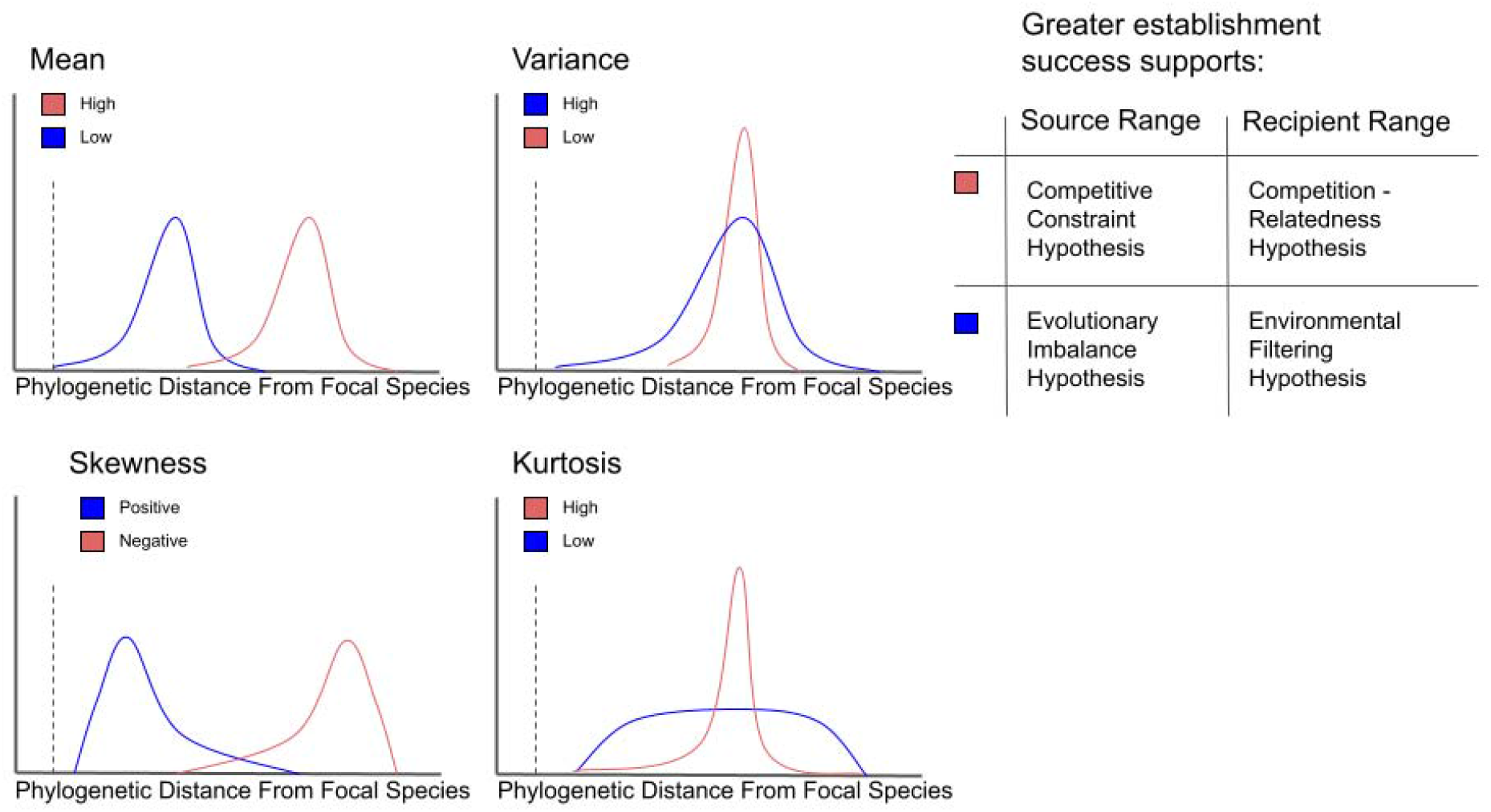
Extending phylogenetic hypotheses of establishment success to include higher moments of phylogenetic distance distributions. Shown are examples of how the shapes of distributions of phylogenetic distances between a community and a single focal species (dotted line) are reflected in their moments. Line colors correspond to support for the different source and recipient range hypotheses (inset, Table 1).

Here we present the first empirical work to examine multiple hypotheses of how both source and recipient range phylogenetic structure will influence establishment success. In order to understand how the inclusion of source range phylogenetic data may alter our conclusions, we revisit the first dataset used to examine the impact of recipient range phylogenetic structure on establishment success in birds (Maitner et al. 2012). This dataset focuses on three highly invaded avifaunas: Florida, Hawaii and New Zealand. In addition to the commonly used metrics of phylogenetic diversity (PD), phylogenetic nearest neighbor distance (NND), and mean phylogenetic distance between species (MPD), we also include metrics based on the higher moments of the distributions of distances between species: variance in phylogenetic distances (VPD), skewness in phylogenetic distances (SPD), and kurtosis in phylogenetic distances (KPD) between species. Finally, we include additional covariates known to explain establishment success from prior studies: propagule pressure (Lockwood et al. 2009) and species traits known to be associated with establishment success (Sol et al. 2012).

## Methods

### Introduction Data

We conducted tests of the alternative source and recipient region hypotheses using data regarding bird introductions to Florida, Hawaii and New Zealand previously used by Maitner *et al. (2012)*. Bird introductions to these regions are relatively common, with relatively high ratios of introduced species to native species (>25%). Published data for both failed and successful introductions are available for all three regions (Long 1981, Moulton et al. 2001, Ornithological Society of New Zealand 2003, Pranty 2004). We analyzed a total of 433 introductions originating from 6 continents (Fig. 1). We followed Maitner *et al.* (2012) in categorizing species as either “successes” or “failures” based on their original classifications in the source materials (Long 1981, Moulton et al. 2001, Ornithological Society of New Zealand 2003), or in the case of Pranty (2004), where a species was listed as breeding and was not listed as having been extirpated. We excluded introduction attempts if: (1) fewer than 5 individuals were introduced (following Moulton et al. 2001); (2) introduction status was listed as uncertain (due to insufficient documentation); or (3) the introduction represented a translocation or reintroduction of a species that was native within the region. Our data set contained 347 species, 79% of which were introduced to one region, 17% to two regions, and 4% to all three regions. These introductions resulted in 30% successes and 70% failures.

**Figure 1.**
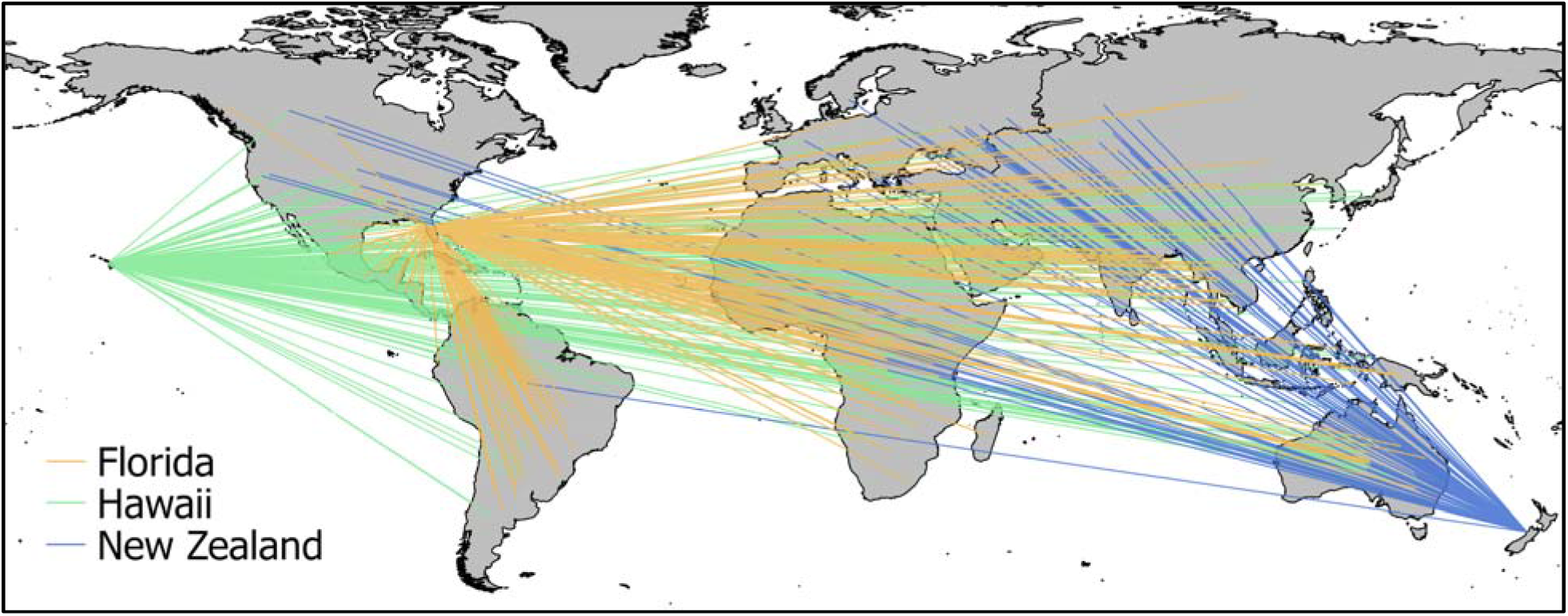
Bird introduction sources for each of the three focal regions of this study (Florida, Hawaii, New Zealand). Lines connect the centroid of the source range of each introduced species with the geographic centroid of the region where it was introduced.

### Source and Recipient Communities

We quantified native and recipient communities using a “phylogenetic field” approach. Phylogenetic fields quantify the phylogenetic relatedness of all the species that co-occur with the focal species of interest (Villalobos et al. 2013), and are a property of species. Quantifying a phylogenetic field provides a framework for linking species co-occurrence patterns to other aspects of a species such as history of invasion, functional traits, or life history (Barnagaud et al. 2014a, Villalobos et al. 2017). To do this, we projected the species range maps to a cylindrical, equal-area projection and rasterized them at a 110 km by 110 km resolution (Hurlbert and Jetz 2007). A species’ source community was defined as the focal species plus all of the species that shared one or more grid cells with the focal species in its native range (Fig. 2). The recipient community was defined as all species that shared one or more grid cells with the focal region (Fig. 2). We delineated communities using a grid cell approach (rather than intersecting range maps) as Hurlbert and Jetz (2007) recommend that range maps are more appropriately used at these relatively coarse scales.

**Figure 2.**
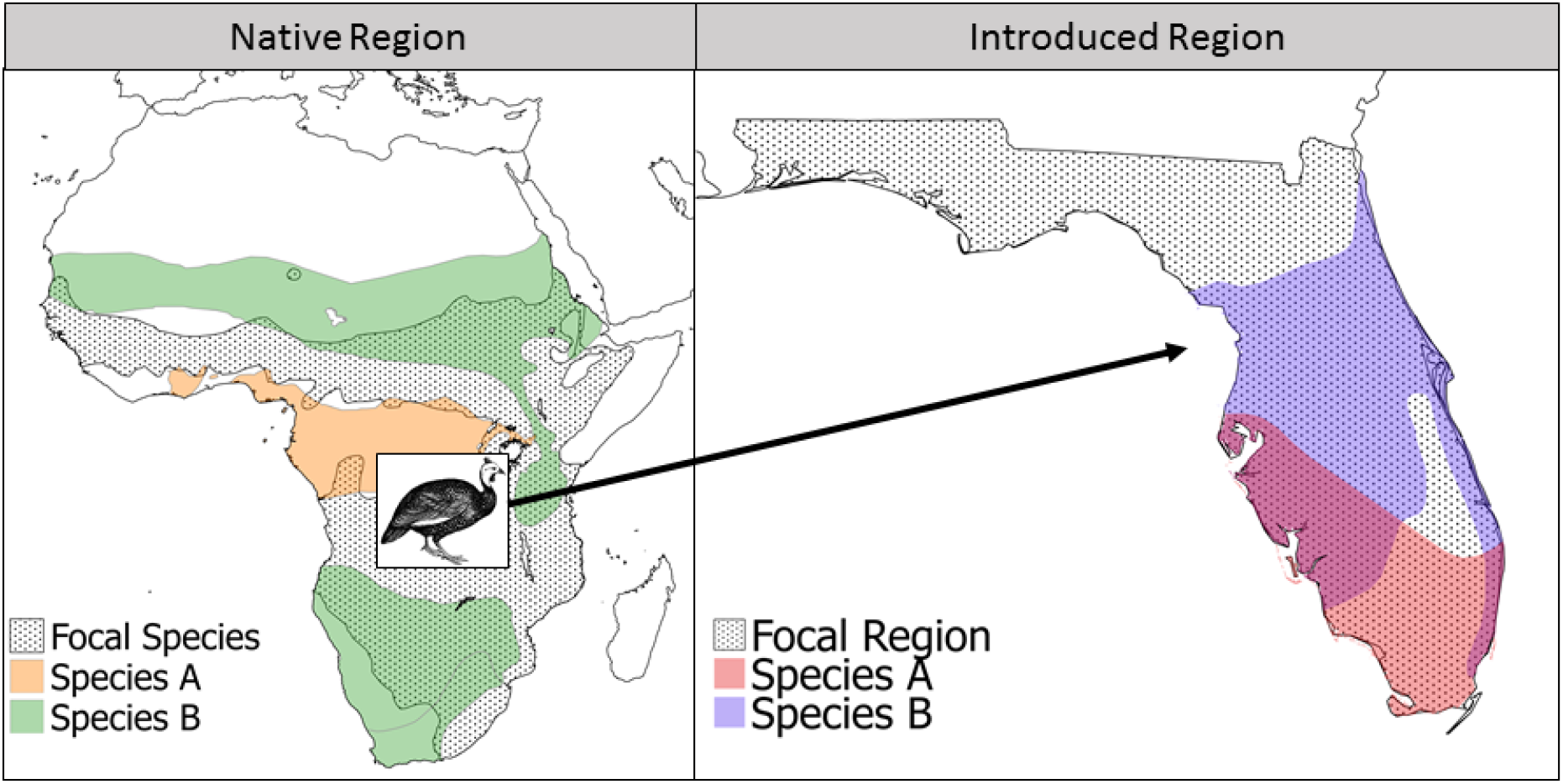
Delineation of native and introduced communities. Within a focal species’ native region, range maps were used to determine which species co-occur with the focal species. Within each introduced region, communities included all species with ranges that intersect the region. Successfully and unsuccessfully introduced species within the region were determined from published sources.

Recipient community data for each region were taken from Birdlife International (BirdLife International 2019) and published sources (Moulton et al. 2001, Ornithological Society of New Zealand 2003, Pranty 2004). Native community data for introduced species were taken from Birdlife International (BirdLife International 2019). In determining native communities we excluded species that were classified as either “introduced” or “vagrant” in that region (BirdLife International 2019). Communities were calculated on the basis of shared grid cells given evidence that this is the finest resolution at which expert range maps can delineate the presence of a species (Hurlbert and Jetz 2007, Kissling et al. 2012). Two species (*Cacatua goffiniana* and *Erithacus komadori*) had ranges so small they were effectively invisible to the rasterization, and were thus omitted from further analyses.

Our definition of the recipient community includes all species with ranges overlapping our focal regions of Florida, Hawaii, and New Zealand and likely overestimates the species that an introduced species encountered. Failed introductions lack a more closely defined recipient community, however, and complete range maps were unavailable for some successful introduced species in our dataset. By including all potential interacting species in each region, we provide the most comprehensive estimate of the recipient community phylogeny, but we expect that this will bias some phylogenetic distance metrics (e.g. nearest neighbor distance will likely be lower; Park et al. 2020).

### Phylogenetic Metrics

For each introduced species we calculated the mean, variance, skewness, and kurtosis of phylogenetic distances between the focal species and all other species in the community (MPD, VPD, SPD, KPD, respectively). Nearest neighbor distance (NND) was calculated as the shortest phylogenetic distance between the focal species and all other species in the community. MPD, VPD, SPD, KPD, and NND within the introduced range were calculated relative to three sets of species: 1) the native community; 2) the native community plus successfully established introduced species; 3) successfully established introduced species only. Calculating the phylogenetic metrics relative to these different communities may show important differences if introduced species are predominantly interacting with a subset of the extant community (e.g. established species; Barnagaud et al. 2014b). Phylogenetic metrics were calculated using the Jetz et al. (2012) phylogenies derived from the Hackett et al. (2008) backbone, which was the most complete avian phylogeny available at the time of these analyses. Native range size was calculated as the number of grid cells that a species’ range intersected. For each of the questions detailed below, introduction success was predicted using linear models, and model fits were compared via Akaike information criterion (AIC).

The Jetz et al. (2012) phylogenies are a pseudo-posterior distribution of phylogenies constructed by grafting species without genetic information within their clade in a manner consistent with taxonomy and inferred branching times (Jetz et al. 2012). These phylogenies are among the best available bird phylogenies and are in frequent use (e.g. Jetz et al. 2014, Freeman et al. 2019, Montaño-Centellas et al. 2020). For community phylogenetic metrics, the use of commonly-available phylogenies is supported, as the results are strongly correlated with results using the more time-consuming approach of generating a purpose-built phylogeny (Li et al. 2019). Nonetheless, to account for potential phylogenetic biases and uncertainty, each phylogenetic metric (PD, NND, MPD, VPD, SPD, KPD) was calculated as the mean value of that metric derived from a random sample of 100 phylogenies from the complete set of 10,000 phylogenies. Across the 100 phylogenies used, the standard error in cophenetic distance was of a relatively small magnitude (1.09 million years) relative to the mean cophenetic distance (159.16 million years).

Phylogenetic diversity was calculated using the function “pd” in the R package *picante* (Kembel et al. 2010). MPD, VPD, SPD, KPD, and NND were calculated by extracting the mean, variance, skewness, kurtosis, and minimum phylogenetic distances (respectively) between a focal species and the rest of the community using the function “cophenetic” in the R package *stats (R Core Team 2020)*.

### Statistical analyses

All analyses were conducted in R version 3.6.3 (R Core Team 2020) using the package package *lme4* (Bates et al. 2015) for non-phylogenetically corrected models and *phyr* (Li et al. 2020) for phylogenetically-corrected models. Model fits were compared via Akaike information criterion (AIC). The scales of the predictor variables differed by orders of magnitude, and so all variables were standardized to a mean of zero and standard deviation of 1 prior to analysis using the function “scale” in R (R Core Team 2020).

#### Comparison of phylogenetic metrics

We compared the relative predictive ability of six native range phylogenetic metrics (PD, NND, MPD, VPD, SPD, KPD) by comparing the fits of generalized linear models. The models contained the source region phylogenetic metric, source region range size, and source region species richness as predictor variables, region of introduction (Hawaii, Florida, New Zealand) as a random effect and introduction success as a binary response (successful vs failed) variable with a logit link. We also compared these models with an equivalent set of phylogenetically corrected models. The predictor variable native range size was significantly correlated with PD (Pearson correlation correlation = 0.38, p < 0.05), marginally correlated with NND (r = −0.09, p = 0.054), and not significantly correlated with MPD, VPD, SPD, or KPD (*p* > 0.05). Species richness was strongly, and significantly correlated with PD (r = 0.99, *p* < 0.05), significantly correlated with NND (r = −0.10, p < 0.05) and VPD (r = −0.12, p < 0.05), and not significantly correlated with MPD, SPD, or KPD (p > 0.05). Due to the strong correlation between species richness and PD, we excluded species richness from models where PD was the response variable.

#### Comparison of competing hypotheses for introduction success

We compared the fits of phylogenetically corrected models containing 1) source community phylogenetic metrics only (i.e. tests of the Evolutionary Imbalance Hypothesis vs. the Competitive Constraint Hypothesis vs. the Universal Trade-Off Hypothesis); 2) recipient community metrics (i.e. tests of Environmental Filtering vs. Competition-Relatedness); and 3) both source and recipient community metrics. Metrics included PD, NND, MPD, VPD, SPD, and KPD. All models included native range size as a covariate, region of introduction as a random effect, and establishment success as the binary response variable with a logit link. All models contained source region species richness except one containing PD, as PD was strongly and significantly correlated with species richness. To account for phylogenetic uncertainty, we fit model models using 100 randomly chosen phylogenies and took the mean values of each model coefficient. We also fit equivalent non-phylogenetically corrected models for comparison purposes using the function “glmer” in the *lme4* R package (Bates et al. 2015).

#### Comparison of phylogenetic metrics with propagule pressure and traits

To compare the predictive power of phylogenetic metrics with that of propagule pressure and species’ traits known to be related to establishment success, we conducted additional analyses on a subset of our data for which data on propagule pressure and other traits were available. Traits included in models were those identified by Sol *et al.* (2012) in their best-fit model and included body mass, the residuals of brain mass against body mass, the value of a brood relative to expected lifetime reproductive output, and habitat generalism (the number of habitat types used by a species). Propagule pressure metrics included both the number of individuals and number of introduction events. We included our best-fitting phylogenetic metrics, source MPD and recipient community MPD (relative to native species only; Table SI 2). We used phylogenetic generalized linear mixed-effect models to test hypotheses based on 1) Propagule pressure only; 2) Propagule pressure and MPD metrics; and 3) Propagule pressure and species’ traits. The “propagule pressure only” model included the number of introduction events and number of individuals introduced as fixed effects. The “propagule pressure and MPD model” additionally included source and recipient community MPD as predictor variables. The “propagule pressure and traits” model included body mass, brain mass residuals, brood value, and habitat generalism in addition to the number of events and individuals as predictor variables. All models included establishment success as a binary response variable with a logit link and a random effect of site (Hawaii or New Zealand). We calculated partial R^2^ using the function “R2_pred” in the *rr2* package for R (Ives and Li 2018), which calculates partial R^2^ as 1 − var(y − y_full_)/var(y − y_reduced_), where y_full_ is the full model and y_reduced_ is a model containing only the intercept (Ives and Li 2018).

## Results

### Comparison of phylogenetic metrics

In our dataset, establishment success showed a weak, non-significant trend of positive association with PD of the native (source) range (Fig. 3, Supplementary Table S1). In contrast, establishment success showed a significant, positive relationship with source range VPD and SPD, as well as significant, negative relationships with NND, MPD, and KPD (Fig. 3, Table S1). Phylogenetically corrected models were broadly consistent with non-phylogenetically corrected models in both sign and significance, with the exception that SPD was no longer significant after phylogenetic correction (Fig. 3). Of the phylogenetically corrected models, the one containing VPD as a predictor showed the best fit (ΔAIC = 3.9; Table S1). The non-phylogenetically corrected PD model performed the worst, and PD was not significantly related to establishment success (Table S1).

**Figure 3.**
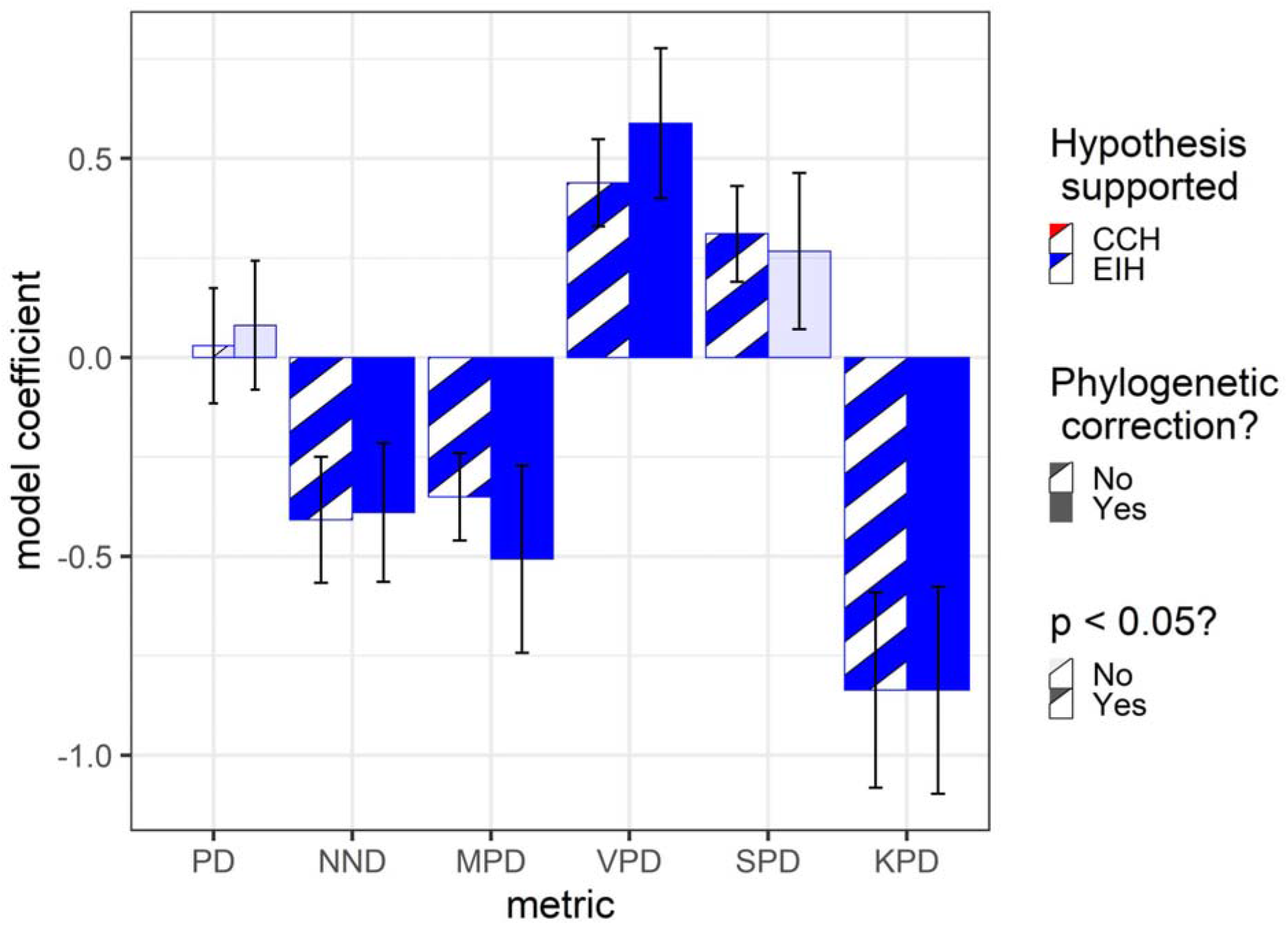
Comparison of phylogenetic metrics in tests of the source range metrics. Shown are estimated model coefficients and standard errors. Lightly filled bars represent model coefficients which weren’t significantly associated with establishment success. Striped bars represent models that weren’t phylogenetically corrected. Blue bars (i.e. all of them) represent models that are consistent with the predictions of the Evolutionary Imbalance Hypothesis. Full models included native region phylogenetic metric (PD, NND, MPD, VPD, SPD, KPD), and species richness and area of the focal species’ native range as predictors, region of introduction as a random effect and introduction success as a binary response variable. Values for the estimates using phylogenetic corrections represent the means of one hundred replicated phylogenies. As PD was strongly correlated with species richness (r = 0.99, p < 0.05), species richness was omitted from that model.

**Figure 4.**
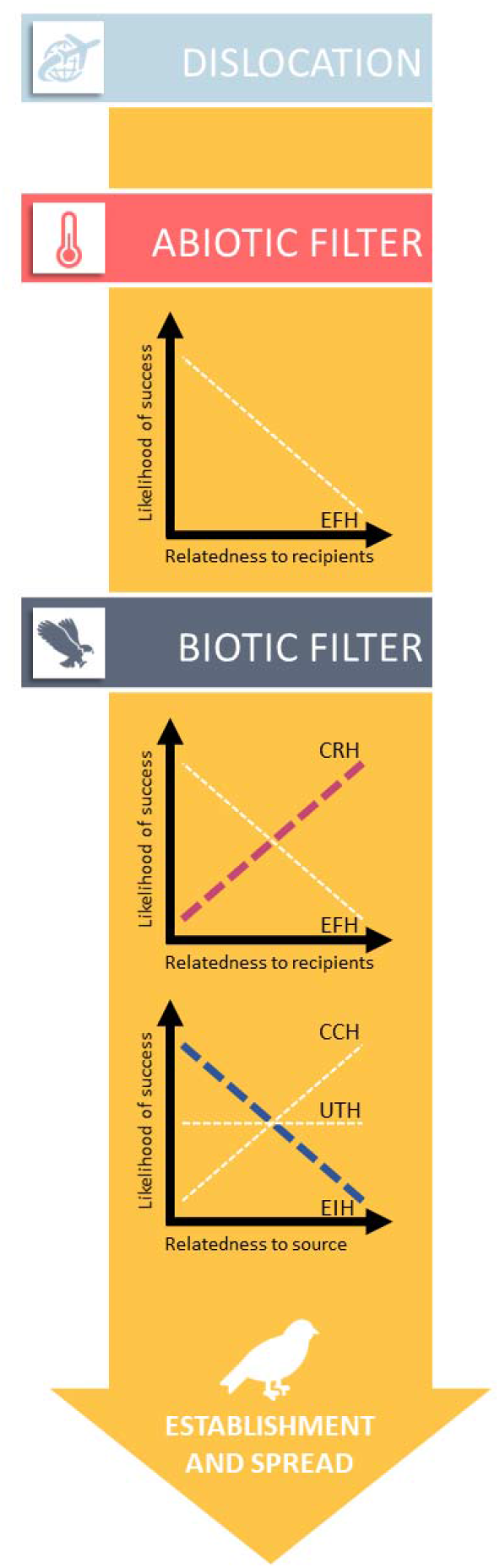
Summary of support for source- and recipient-range hypotheses. Dashed lines reflect hypothesized relationships. Thick, colored lines denote relationships that are supported when including both source- and recipient-range phylogenetic information. The thin, white lines indicate hypotheses that are not supported here. CRH = Competition-Relatedness Hypothesis, EHF = Environmental Filtering Hypothesis, CCH = Competitive Constraint Hypothesis, UTH = Universal Trade-off Hypothesis, EIH = Evolutionary Imbalance Hypothesis.

### Comparison of competing hypotheses for introduction success

Our preferred model (ΔAIC > 2) included a marginally significant (0.1 > p > 0.05) positive effect of source region range size, a non-significant effect of source region species richness, a significant (p < 0.01) negative effect of source region MPD, and a significant (p < 0.01) positive effect of recipient region MPD relative to native species only (Table SI 2). We also found that across all of our models the effect of source region phylogenetic metrics was consistent in sign, with NND, MPD, and KPD always showing a negative relationship with establishment success, and VPD, and SPD always showing a positive relationship with establishment success. In contrast, most recipient region phylogenetic metrics (with the exception of NND, which always showed a negative relationship with establishment success) showed inconsistent relationships with establishment success that changed in direction depending on circumscription of the recipient species pool (i.e. native, established, or native and established species) and whether source region phylogenetic structure was included Source region range size always showed a positive association with establishment success that was either significant (p < 0.05) or marginally significant (0.1 > p > 0.05). Source region species richness was never significantly related to establishment success, and varied in sign.

The non-phylogenetically corrected models (Table SI 3) were broadly consistent with the phylogenetically-corrected models: the best-performing model again included a significant, negative effect of source-range MPD, and a significant, positive effect of recipient-range MPD (relative to the native community). As with the phylogenetically-corrected models, source-region NND, MPD and KPD showed negative relationships with establishment success, while source region VPD and SPD showed positive relationships (Table SI 3), and recipient community metrics were sensitive to the additional terms included and the circumscription of the recipient community. However, there were some important differences, including model terms that differed in sign and/or significance between the models. For example, recipient-range MPD calculated to the native community only shows a positive, significant effect when correcting for phylogeny, but a negative, non-significant effect when failing to correct for phylogeny.

### Comparison of phylogenetic metrics with propagule pressure and traits

From our initial dataset consisting of 433 observations, we retained 112 observations for which trait and propagule pressure data were available for introductions into Hawaii and New Zealand. In our “propagule pressure only” model (mean partial r^2^ = 0.428, SE = 0.001), we found a significant (p < 0.05), positive effect of the number of introduction events on establishment success, but did not find a significant effect of the number of individuals introduced. Our model including community phylogenetic structure metrics in addition to propagule pressure offered an improved fit relative to the propagule pressure only model (ΔAIC = 3.41, mean partial r^2^ = 0.417, SE = 0.001), and included a significant (p < 0.05), negative effect of source community MPD and a significant (p < 0.05), positive effect of the number of introduction events on establishment success, while recipient region MPD and the number of individuals introduced were not significantly related to establishment success. Our propagule pressure and traits model provided the best fit of the three models examined (ΔAIC > 93, mean partial r^2^ 0.576, SE = 0.001), and included a significant (p < 0.05), positive effects of both habitat generalism and the number of introduction events on establishment success, while body mass, residual brain mass, brood value, and the number of individuals introduced were not significantly related to establishment success.

## Discussion

Negative relationships between establishment success and source MPD and NND, and a positive relationship with source SPD (although SPD was only significant prior to phylogenetic correction), indicate that species with more close relatives (low values of MPD, NND, high values of SPD) in their native range had higher introduction success elsewhere. A positive relationship of source VPD and a negative relationship of source KPD with establishment success indicate that species which co-occur with species representing a greater diversity of phylogenetic distances had higher introduction success. Thus our initial findings were consistent with the predictions of the Evolutionary Imbalance Hypothesis, and there was a clear association between species that come from closely-related and/or diverse communities and increased establishment success (Fig. 3).

Consistent with predictions of the Evolutionary Imbalance Hypothesis, the overall best phylogenetic model included a significant, negative relationship between introduction success and source range MPD, and a significant, positive effect of recipient range MPD calculated relative to the native community. In other words, species originating in regions with communities containing relatively closely-related species that are then introduced into regions containing relatively distantly-related species are more likely to become established. The next-best model was a substantially poorer fit (ΔAIC = 3.26), and included a significant, positive relationship between establishment success and source VPD, and a significant, negative effect of recipient range VPD relative to the native community(Table SI 2). In other words, species coming from regions where they span a diverse range of phylogenetic distances to the community are most likely to succeed when introduced into a community where they encounter a smaller variety of phylogenetic distances.

Importantly, we found that including source range phylogenetic structure in models, changing the circumscription of the recipient community (e.g. including established and/or native species), and utilizing phylogenetic corrections can change the sign and significance of model terms. Including native range MPD in the non-phylogenetically corrected model caused a change in both the direction and significance of the effect of recipient range MPD, from a (nonsignificant) negative relationship as predicted by Environmental Filtering (similar to the findings of Maitner et al. 2012), to a significant positive relationship as predicted by Competition-Relatedness (Table SI 3). However, this was not true when employing phylogenetic corrections (Table SI 2). Therefore the decision of whether (or not) to incorporate both source and recipient region phylogenetic structure, or to include phylogenetic corrections in an analysis can fundamentally change the results, and may warrant re-analysis of previously published work.

Adding phylogenetic structure (source and recipient community MPD) to models containing propagule pressure alone improved the fit (ΔAIC = 3.41), however the the effect of recipient region phylogenetic structure was not significantly associated with establishment success, unlike in the model containing only phylogenetic structure metrics. One potential explanation for these differences between the models with propagule pressure and those without them is that the subset of introductions with information about propagule pressure differs from our full dataset, and might have a weaker relationship with recipient range phylogenetic structure. We tested for this possibility and indeed found that source range MPD -- but not recipient range MPD -- was significantly related to establishment success in a model that utilized only the 112 observations with information about propagule pressure. This suggests that within this subset of data, source region MPD is indeed a stronger predictor of introduction success than recipient region MPD.

These findings support the well-established importance of propagule pressure in predicting introduction success (Holle and Simberloff 2005, van Wilgen and Richardson 2012, Blackburn et al. 2015), but also indicate that phylogenetic metrics can add significant explanatory power to these models when trait information is lacking. Detailed trait data were yet more powerful for predicting establishment in our dataset, but these types of data are not always readily available and there is a persistent desire to use phylogenetic structure as a proxy for ecologically relevant traits (Van Wilgen and Richardson 2011, Violle et al. 2011, Ma et al. 2016).

Previous tests of recipient community phylogenetic structure on the establishment success of birds have found that species that are more closely-related to the recipient community were more likely to successfully establish (consistent with Environmental Filtering; Maitner et al. 2012, Baiser et al. 2017) and that the NND of the recipient community was a better predictor of establishment success than MPD (Maitner et al. 2012). We find similar patterns when only considering the recipient community NND in our dataset (Tables 2, 3). In contrast, our best-fit model provides support for the opposite conclusion when accounting for native region phylogenetic structure, that species that are more distantly-related to the recipient community are more likely to successfully establish. Intriguingly, a recent ecological study of established bird species in New Zealand found that native and introduced species tend to occupy different habitats and have relatively low overlap in functional trait space (Barnagaud et al. 2014b), suggesting differentiation rather than environmental filtering is structuring avian establishment in this region, as indicated by our model incorporating the Evolutionary Imbalance Hypothesis here.

The results presented here utilize data from three distinct regions which were selected for their high availability of data for both successful and failed introductions. However, it is possible that these regions may not be broadly representative. Both Hawaii and New Zealand are islands, while Florida is a climatic “island”, separated from other tropical and subtropical regions by both water and temperate climates (Hardin 2007). It has been suggested that islands may be easier to invade (Elton 1958), however, evidence suggests this may not be the case for birds (Sol 2000). The importance of biotic interactions in these introductions may also not be broadly representative, as both Hawaii and New Zealand have had many extinction events following the arrival of humans (Blackburn et al. 2004), and it has also been suggested that the tropical and neotropical regions of Florida may also be relatively depauperate owing to their climatic isolation (Hardin 2007). We recommend caution in the interpretation of our findings until we can establish whether they are broadly representative of introductions more generally.

## Conclusions

Recent years have seen an active interest in revisiting and testing Darwin’s (1859) ideas regarding phylogenetic patterns in introduction success. There have been many tests of the Competition-Relatedness (‘Naturalization’) and Environmental Filtering hypotheses in particular e.g. (Daehler 2001, Strauss et al. 2006, Diez et al. 2008, Thuiller et al. 2010, Van Wilgen and Richardson 2011, Violle et al. 2011, Maitner et al. 2012, van Wilgen and Richardson 2012, Park and Potter 2015a, b, Li et al. 2015, Ma et al. 2016, Park and Razafindratsima 2019), but these have culminated in an often frustrating lack of generality (Thuiller et al. 2010, Ma et al. 2016, Cadotte et al. 2018, Park et al. 2020). These hypotheses focus on phylogenetic relationships with recipient communities, but Darwin and others have also considered the potential for the evolutionary history of introduced species in their source communities to influence introduction success. The formalization of multiple source-region hypotheses (reviewed in Maitner et al. 2021) has emphasized the need to test these ideas, and raised the possibility that incorporating both source and recipient community features could resolve general patterns in the ecophylogenetics of introduced species.

Our case study examined the success of birds introduced into three well-described avian communities. Despite previous support for the Environmental Filtering Hypothesis in these introductions (Maitner et al. 2012), here we found support for the Competition-Relatedness Hypothesis once we accounted for the phylogenetic structure of species’ source regions. We also found consistent support for the Evolutionary Imbalance Hypothesis. These results indicate that accounting for both source and recipient communities could both provide important insights from the source region relationships, and improve the ability to detect competitive effects within recipient communities at a regional scale.

While there has been relatively little work to date focusing on source range hypotheses, these have received indications of support from a combination of empirical (Tilman 2011, Fridley and Sax 2013), theoretical (Tilman 2011, Wilson 2014), and experimental studies (Korona 1996, Meyer and Kassen 2007). We note that there are a number of important considerations that will arise when testing these hypotheses in a phylogenetic context and interpreting the results. These include the choice of phylogenetic metric, whether to include native or established species (or both) in the recipient community, the spatial scale and resolution modeled, and the inclusion of other explanatory factors such as range size, propagule pressure, or species’ traits. It may be especially important to recognize that, while including species traits or propagule pressure can improve predictive power (van Wilgen and Richardson 2012, this study), these explanatory factors may also covary with phylogenetic patterns, particularly where there are large ecological and propagule pressure differences among major clades (Westoby et al. 1995). A particularly pressing issue is the degree to which any best fit model has substantial explanatory power in predicting introduction success (Mac Nally et al. 2017). These issues identify the need for analyses exploring how phylogenetic metrics, species’ traits, propagule pressure, and community circumscription, within both source and recipient communities, impact predictions of invasion success across scales, as well as theoretical and experimental studies to refine predictions and test specific mechanisms likely to generate observed patterns

## Acknowledgements

KMD was supported by USDA #2015-67013-23000, NSF #1550838, and NSF #1750280. BJE and BSM were supported by National Science Foundation award DEB 1457812 and a Conservation International SPARC award to BJE. BJE was also partially supported by a Visiting Professorship Grant from the Leverhulme Trust, UK and an Oxford Martin School Fellowship. We thank PC and JF for helpful comments on previous drafts. Any opinions, findings, and conclusions or recommendations expressed in this material are those of the authors and do not necessarily reflect the views of the National Science Foundation.

## Supporting Information

**Table S1.** Comparison of native range metrics. Statistical models contained the source region phylogenetic metric, source region range size, and source region species richness as predictor variables, region of introduction (Hawaii, Florida, New Zealand) as a random effect and introduction success as a binary response (successful vs failed) variable. We also compared these models with an equivalent set of phylogenetically corrected models. Due to the strong correlation between species richness and PD, we excluded species richness from models where PD was the response variable.

| metric | estimate | s.e. | p value | phylogenetically corrected? | AIC |
| --- | --- | --- | --- | --- | --- |
| VPD | 0.43867 | 0.109662 | 6.33E-05 | No | 500.7958 |
| VPD | 0.588434 | 0.188717 | 0.001833 | Yes | 501.317 |
| KPD | -0.83636 | 0.245605 | 0.000661 | No | 502.7228 |
| KPD | -0.83644 | 0.260522 | 0.001338 | Yes | 505.0492 |
| MPD | -0.35087 | 0.109504 | 0.001355 | No | 506.2977 |
| MPD | -0.50684 | 0.235319 | 0.031518 | Yes | 506.3459 |
| NND | -0.39012 | 0.174652 | 0.025581 | Yes | 507.1687 |
| NND | -0.4082 | 0.158132 | 0.009841 | No | 508.511 |
| SPD | 0.310473 | 0.120438 | 0.009942 | No | 509.667 |
| PD | 0.080609 | 0.161769 | 0.618554 | Yes | 509.8142 |
| SPD | 0.26718 | 0.196375 | 0.176325 | Yes | 510.3789 |
| PD | 0.028877 | 0.144816 | 0.841945 | No | 514.92 |

**Table SI 2.**
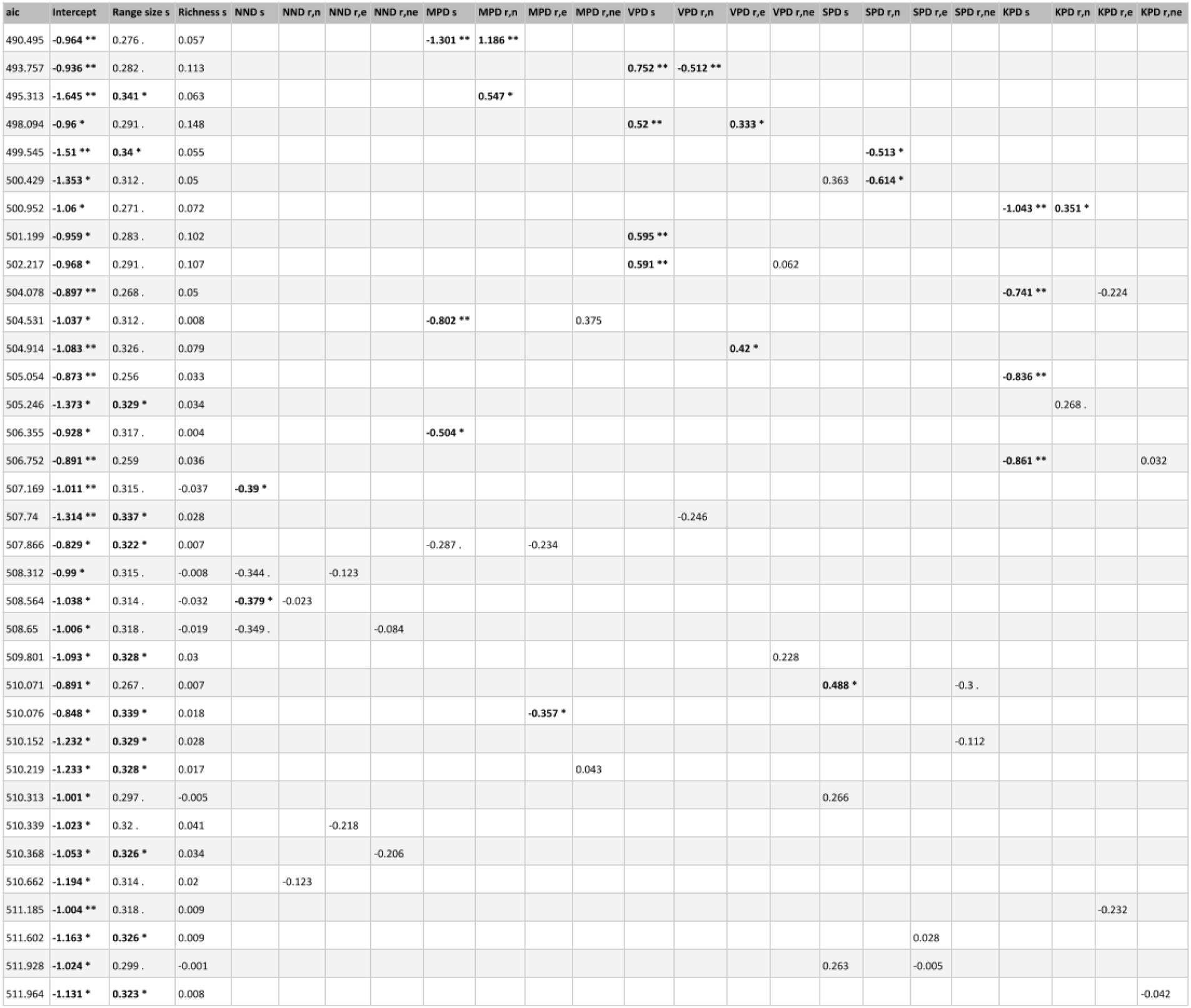
Phylogenetically-corrected generalized linear model summary table. Coefficients shown are means taken across 100 replicated phylogenies. Suffixes ‘r’ and ‘s’ denote the to which the metrics refer: ‘s’ denotes the source region and ‘r’ denotes the recipient community. Suffixes ‘n’ and ‘e’ denote how the recipient communities were circumscribed: ‘n’ included only native species; ‘e’ included only established species, and ‘ne’ included both native and established species. Models are sorted in order of increasing AIC. The symbols “.”, “*”, and “**” denote 0.1 > p > 0.05, 0.05 > p > 0.01, and p < 0.01, respectively. Significant values (p < 0.05) are in bold.

**Table SI 3.**
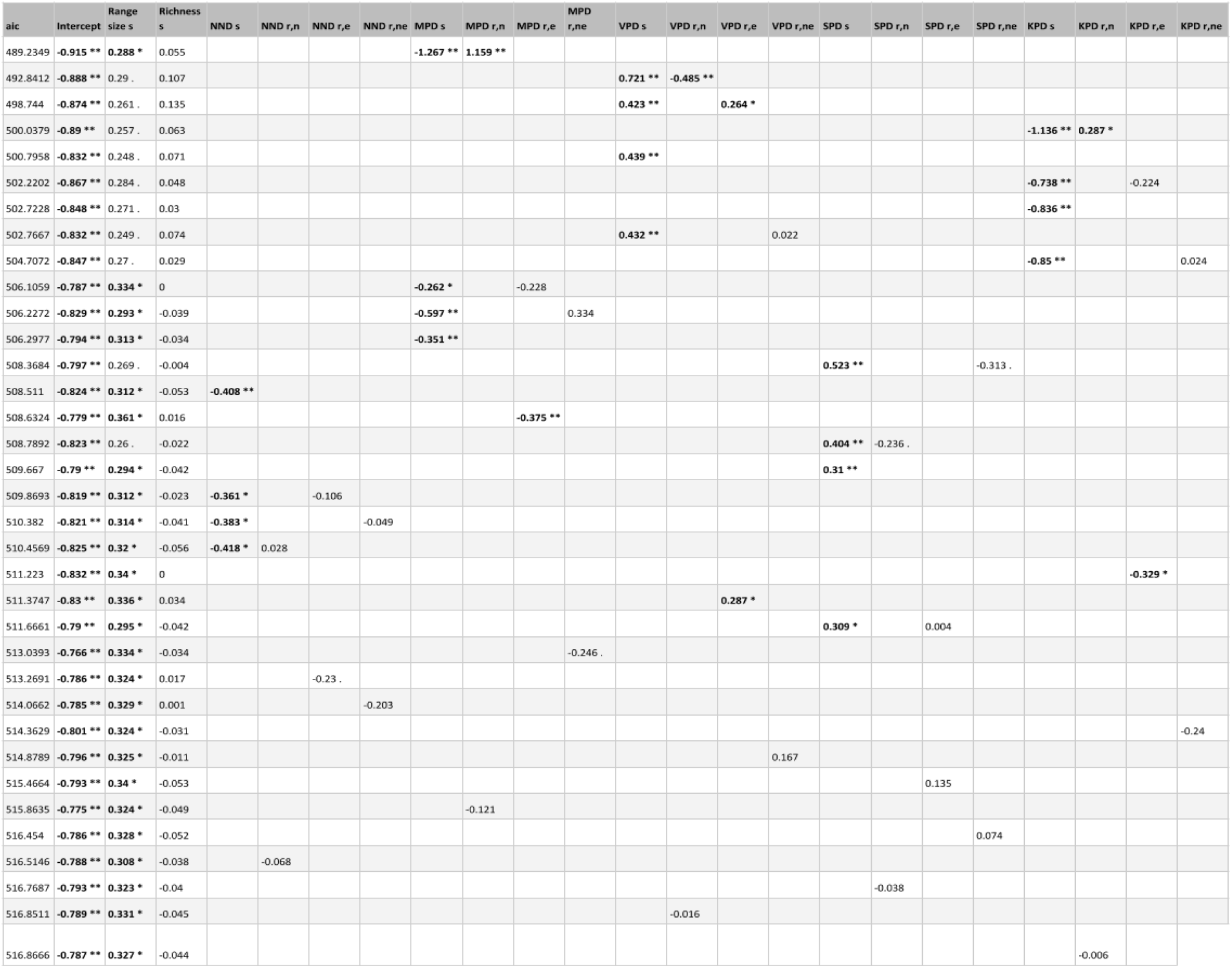
Non-phylogenetically-corrected generalized linear model summary table. Suffixes ‘r’ and ‘s’ denote the to which the metrics refer: ‘s’ denotes the source region and ‘r’ denotes the recipient community. Suffixes ‘n’ and ‘e’ denote how the recipient communities were circumscribed: ‘n’ included only native species; ‘e’ included only established species, and ‘ne’ included both native and established species. Models are sorted in order of increasing AIC. The symbols “.”, “*”, and “**” denote 0.1 > p > 0.05, 0.05 > p > 0.01, and p < 0.01, respectively. Significant values (p < 0.05) are in bold.

## Statement of authorship

BSM and KMD conceived the study, BSM, BJE, and KMD refined and contributed to its design. BSM performed the research. BSM and KMD analyzed the data and drafted the manuscript. BSM, DSP, BJE, and KMD edited the final manuscript.

## Credit statements

**BSM:** Conceptualization(Lead), Formal Analysis (Lead), Visualization (Lead), Writing - Original Draft (Lead), Writing - Review & Editing

**DSP:** Writing - Review & Editing, Visualization (Supporting)

**BJE:** Funding acquisition (Lead), Supervision (Supporting), Writing - Review & Editing

**KMD:** Funding acquisition(Supporting), Conceptualization(Co-Lead), Supervision (Lead), Formal Analysis (Supporting), Writing - Original Draft (Supporting), Writing - Review & Editing

## Data accessibility statement

If this manuscript is accepted for publication, the supporting data will be archived in an appropriate public repository.

